# *Butyribacter intestini* gen. nov., sp. nov., a butyric acid-producing bacterium of the family *Lachnospiraceae* isolated from the human faeces, and reclassification of *Acetivibrio ethanolgignens* as *Acetanaerobacter ethanolgignens* gen. nov., comb. nov

**DOI:** 10.1101/2020.09.01.276766

**Authors:** Wenbin Xue, Xiaoqian Lin, Mei Lv, Guangwen Luo, Ying Dai, Haipeng Sun, Liang Xiao, Yuanqiang Zou

## Abstract

A novel, non-motile, Gram-stain-positive, non-spore-forming, obligate anaerobic bacterium, designated strain TF01-11^T^, was isolated from human faeces. The isolate was characterized by phylogenetic and phenotypic properties, as well as by determination of its whole genome sequence. The growth temperature and pH ranges were 30–42 °C and 6.0–8.5, respectively. The end products of glucose fermentation were butyric acid and a small amount of acetic acid. The genome was estimated to be 3.61Mbp with G+C content of 36.79 mol%. Genes related to biosynthesis of diaminopimelic acid, polar lipids, polyamines, teichoic and lipoteichoic acids were present. The predominant fatty acids were C_16:0_ (37.9 %), C_14:0_ (16.4 %), C_13:0_ OH/iso-C_15:1_ H (11.1 %) and C_18:1_ *ω*9*c* (10.6 %). Phylogenetic analyses based on 16S rRNA gene sequences, the isolate was a member of family *Lachnospiraceae*, with the highest sequence similarity to the type strain of *Roseburia intestinalis* DSM 14610^T^ at 92.18 % followed by *Acetivibrio ethanolgignens* ATCC 33324^T^ at 91.99 %. The average nucleotide identity (ANI) calculated for the genomes between strain TF01-11^T^ and these closest relatives were 70.5 % and 68.1 %. Based on results of phenotypic characteristics and genotypic properties presented in this study, strain TF01-11^T^ represent a novel species in a new genus, for which the name *Butyribacter intestini* gen. nov., sp. nov. is proposed. The type strain of the type species is TF01-11^T^ (CGMCC 1.5203^T^ = CGMCC 10984^T^ = DSM 105140^T^). In addition, *Acetivibrio ethanolgignens* is proposed to be reclassified as *Acetanaerobacter ethanolgignens* gen. nov., comb. nov.

## Introduction

The human intestine is colonized by a large number of microbial communities, which is 10 times higher than the total cells in the human body [1]. In total there are more than 4000 different species in the gut (Almeida *et al.*, 2020). Most of the intestinal microbiota species are obligate anaerobes [2]. These gut microbiota play a key role involving nutrition extraction [3], host metabolic [4–6], prevention against pathogens [7] and immune regulation [8, 9]. Current evidence also suggests that the gut microbiota can be considered an environmental factor in development of disease, including obesity [10, 11], diabetes [11–13], inflammatory bowel disease [14, 15] and colorectal cancer [16–18]. Member of the family *Lachnospiraceae* was one of the most abundant groups within the *Firmicutes* [19, 20]. The production of butyric acid, a short-chain fatty acid, links with *Lachnospiraceae* has potentially beneficial effects on the host [21–23]. Butyrate is considered as one beneficial metabolites, which serves as the major energy source of intestinal epithelial cells and has anti-inflammatory properties [24, 25].

In this study, we report on the taxonomic characterization of a new butyrate-producing bacterial strain, TF01-11^T^, which was isolated from the faeces of a 17-year-old Chinese female. On the basis of the phenotypic, chemotaxonomic, genotypic and phylogenetic data, strain TF01-11^T^ represents a novel genus in the family *Lachnospiraceae*, with the proposed name, *Butyribacter intestini* gen. nov., sp. nov. Furthermore, we suggest reclassification of *Acetivibrio ethanolgignens* to *Acetanaerobacter ethanolgignens* gen. nov., comb. nov.

## Material and methods

### Sample collection and bacteria isolation

The fresh faeces sample was transferred immediately to anaerobic box (Bactron Anaerobic Chamber, Bactron□-2, shellab, USA) and suspended in 0.1 M PBS (pH 7.0) after collected from a a 17-year-old Chinese female living in Shenzhen. The faeces sample was tenfold diluted and spread-plated onto peptone-yeast extract-glucose (PYG) plates and incubated under anaerobic condition (contained 90 % nitrogen, 5 % hydrogen and 5 % carbon dioxide, by vol.) at 37 °C for 3 days. Single colonies were picked and streaked onto PYG agar until a pure culture was obtained according the method previously [26]. The strain was maintained in glycerol suspension (20 %, v/v) at −80 °C and preserved by lyophilization at 4 °C.

### Phenotypical characterization

Cellular morphology of strain TF01-11^T^ was examined using phase contrast microscopy (Olympus BX51, Japan) by using cells grown in prereduced anaerobically sterilized PYG broth at 37 °C for 24 h. The Gram reaction and spore formation were performed by staining using Gram stain kit (Solarbio) and spore stain kit (Solarbio) according to the manufacturer’s instructions. Motility of cells grown in PYG broth was examined by phase-contrast microscopy. All growth experiments, described below, were evaluated using the PYG medium in two replicates (1 %, v/v, inoculum) and recorded by measuring the OD_600_ of the cultures after 24, 48 h and 7 d, respecitively. The temperature range for growth at 4, 10, 15, 20, 25, 30, 37, 42, 45 and 50 °C and the pH range for growth was assessed at pH 3.0–10.0 (at interval of 0.5 pH units). Salt tolerance was determined in PYG broth containing 0–6.0 % (w/v) NaCl (at 1.0 % intervals). Biochemical reactions and carbon source utilization were investigated by using the API ZYM, API 20A and API 50CH tests (bioMe’rieux) according to the manufacturer’s instructions. Short-chain fatty acids (SCFA) produced from fermentation in PYG medium was measured by gas chromatograph (GC-2014C, Shimadzu) using capillary columns packed with porapak HP-INNOWax (Cross-Linked PEG, 30 m × 0.25 mm × 0.25 um) and detected with a flame-ionization detector. Column temperature was 220 °C, N_2_ was used as the carrier gas in all analyses.

### Chemotaxonomic analyses

Cells grown for 48 h at 37 °C on PYG agar plates were used for the whole-cell fatty acid and peptidoglycan analyses. Cellular fatty acids (CFA) of strain TF01-11^T^ and related species were extracted, methylated and analysed by GC as described previously (Chen & Dong, 2004). The analysed of peptidoglycan structure was carried as described previously [27].

### Phylogenetic analysis based on 16S rRNA gene

The genomic DNA was extracted from cells grown in PYG broth for 24 h at 37°C and purified using the method described by Drancourt *et al.* [28]. The 16S rRNA gene sequence was amplified by PCR and sequenced as described previously [27]. The sequence obtained was compared with entries in EzBioCloud server [29]. Phylogenetic analysis was performed by using software package MEGA 7 [30]. Sequences of TF01-11^T^ and related type species were aligned and used to construst a phylogenetic tree by the neighbour-joining method [31] and maximum likelihood method using CLUSTAL W [32]. In each case, bootstrap values were calculated based on 1000 replications.

### Whole Genome analysis

The draft whole-genome sequence of strain TF01-11^T^ was performed by using a paired-end strategy with the platform Illumina HiSeq 2000 at BGI-Shenzhen (Shenzhen, China). The paired-end library had an mean insert length of 500 bp. Paired-end de novo genome assembly was performed with SOAPdenovo 2 package [33]. The genome sequence of strain TF01-11^T^ was compared with available genome sequences of representatives of the family *Lachnospiraceae*. All genome sequences were obtained from the GenBank sequence database. Individual coding sequences were annotated using the Rapid Annotation Subsystem Technology (RAST) server [34], freely available at (http://rast.nmpdr.org). DNA base content (mol% G+C) was calculated from the whole genome sequence. The average nucleotide identity (ANI) values were calculated for strain TF01-11^T^ and the most closely related species *R. intestinalis* DSM 14610^T^, *A. ethanolgignens* ATCC 33324^T^, *Lachnospira multipara* DSM 3073^T^ and *Coprococcus eutactus* ATCC 27759^T^.

## Results and discussion

### 16S rRNA gene sequencing and phylogenetic analyses

The almost complete 16S rRNA gene sequence of strain TF01-11^T^ of 1,400 bp was determined. Comparative sequence analysis of strain TF01-11^T^ and validly published names using the EzBioCloud server revealed that the most similar sequences were those of the members of family *Lachnospiraceae* of the phylum *Firmicutes*. The 16S rRNA gene sequence similarity of strain TF01-11^T^ and the closest relatives, *Roseburia intestinalis* DSM 14610^T^ [35] and *Acetivibrio ethanolgignens* ATCC 33324^T^ [36], were 92.18 % and 91.99 % (Table 3), respectively, which was below the ‘lower cut-off window’ of 95 % for differentiation of a new genus [37, 38]. Furthermore, the phylogenetic analysis showed that strain TF01-11^T^, together with the closest relative *A. ethanolgignens* ATCC 33324^T^, formed a separate branch within the family *Lachnospiraceae* (Fig. 1 and Supplementary Fig. S1). Additionally, the type species of the genus *Acetivibrio*, *Acetivibrio cellulolyticus*, is phylogenetically classified in the family *Ruminococcaceae* according to the taxonomy list in LPSN (https://lpsn.dsmz.de/genus/acetivibrio) and shared a low 16S rRNA gene sequence similarity (85.2 %) with *A. ethanolgignens* ATCC 33324^T^. It is obvious that strain *A. ethanolgignens* ATCC 33324^T^ does not cluster together with *Acetivibrio cellulolyticus* in the family *Ruminococcaceae,* but more closely group in the family *Lachnospiraceae.* This result was also confirmed by the maximum-likelihood method (Supplementary Fig. S1), suggesting that *A. ethanolgignens* ATCC 33324^T^ should be reclassified as a new genus of the family *Lachnospiraceae.*

### Whole genome sequencing and G+C content

Sequencing of the genome produced an annotated genome size of approximately 3.61 Mbp. The G+C content of DNA was 36.79 mol% as calculated from the whole-genome sequence (Table 2). The ANI between strain TF01-11^T^ and *R. intestinalis* DSM 14610^T^, *A. ethanolgignens* ATCC 33324^T^, *L. multipara* DSM 3073^T^ and *C. eutactus* ATCC 27759^T^ had a maximum value of 70.5 % (Table 3).

**Table 1.**
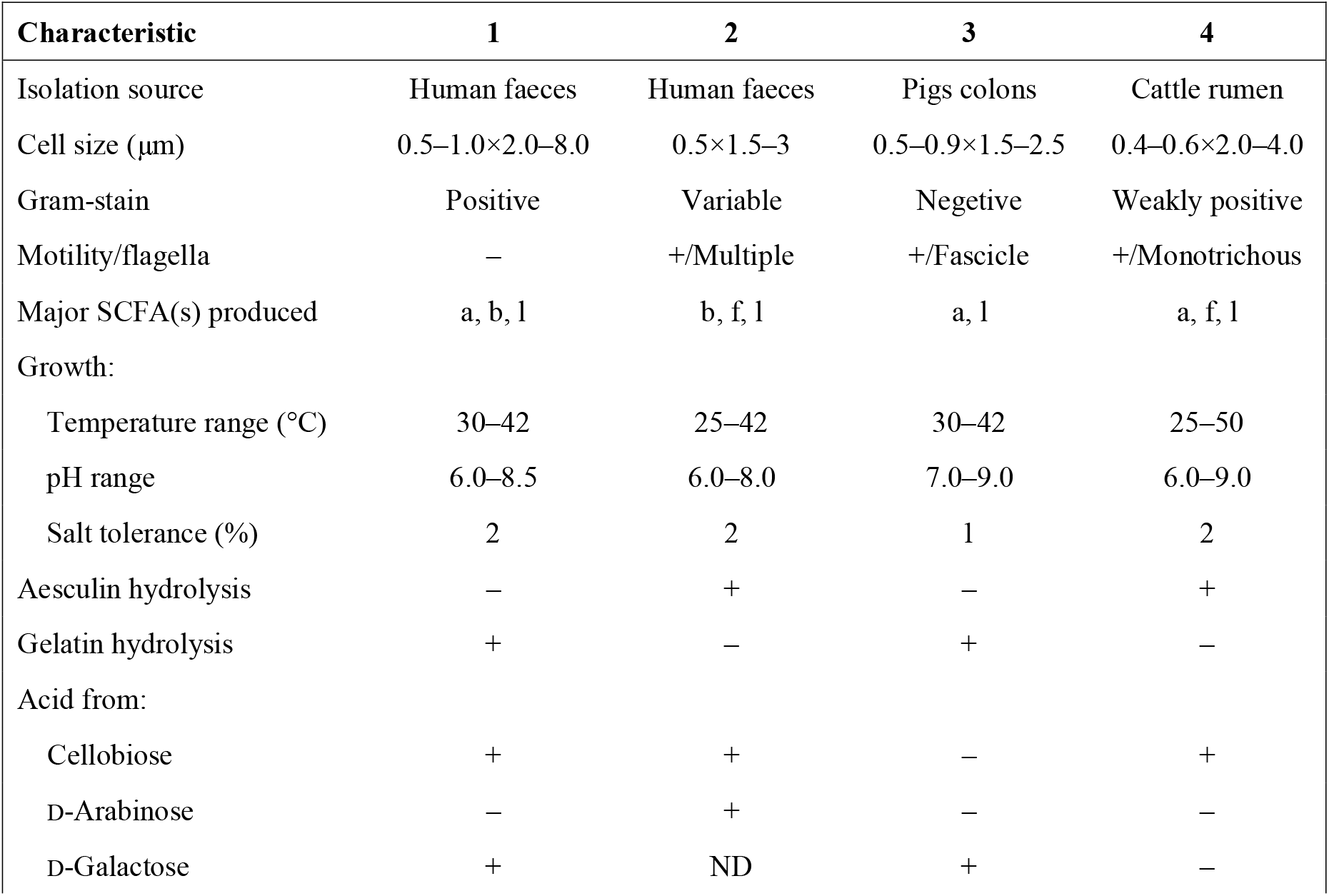
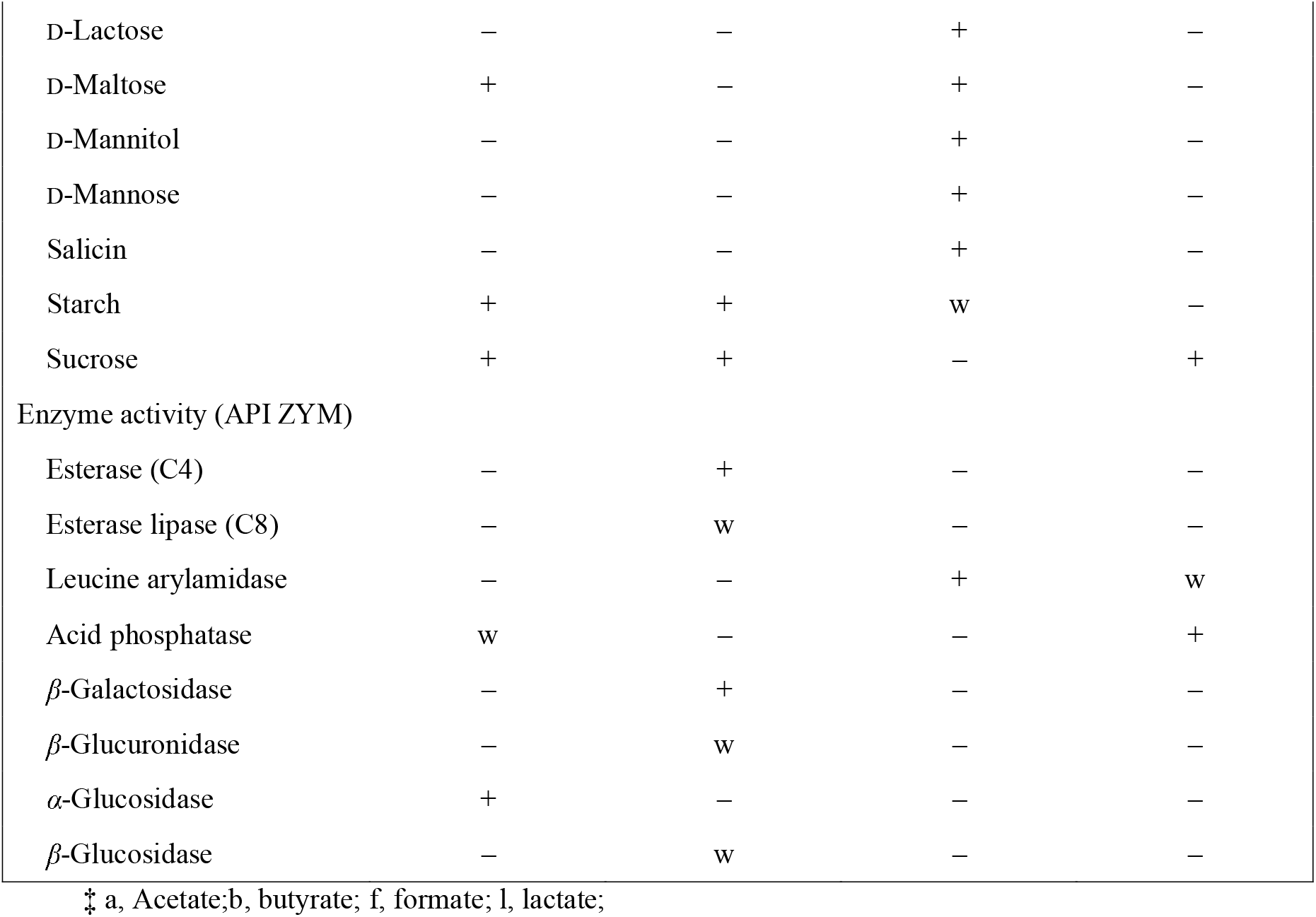
Differential phenotypic features among TF01-11^T^ and the type species of phylognetically closely related members of family *Lachnospiraceae.* Strains: 1, TF01-11^T^; 2, *R. intestinalis* DSM 14610^T^; 3, *A. ethanolgignens* ATCC 33324^T^; 4, *L. multipara* DSM 3073^T^. Data were from Duncan *et al.* (2006), Robinson & Ritchie (1981), Bryant (1986) and this study. +, Positive; w, weakly positive reaction; −, negative; ND, no data available; SCFA, short-chain fatty acid.

**Table 2.**
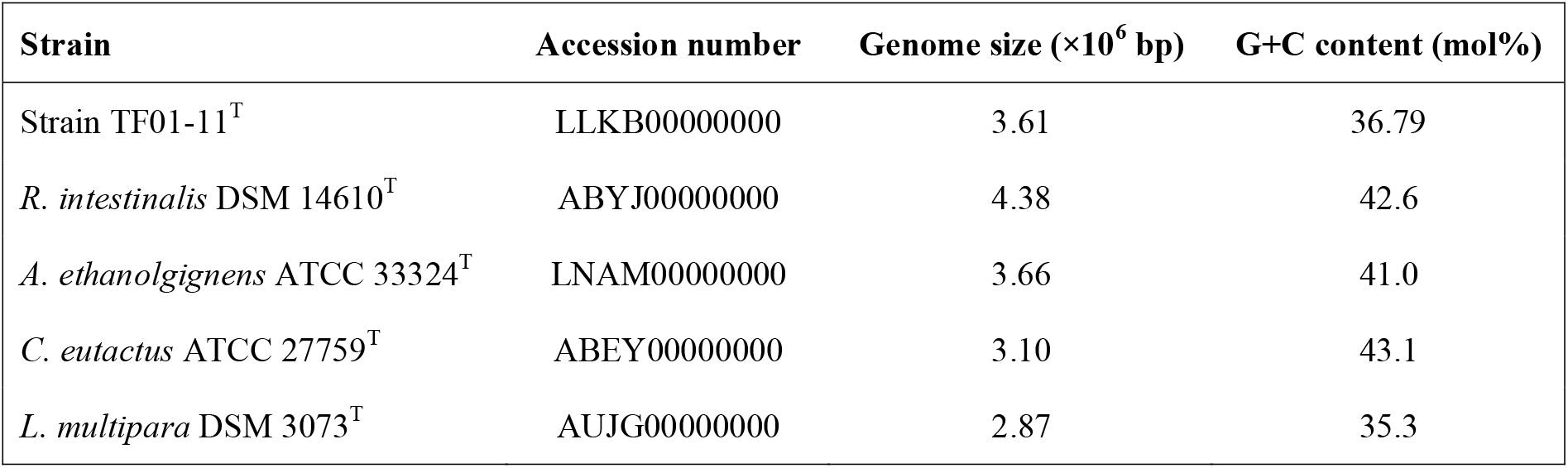
Genome size and DNA G+C content of Strain TF01-11^T^ compared with three closely related isolates of the family *Lachnospiraceae*. The data of *R. intestinalis* DSM 14610^T^, *A. ethanolgignens* ATCC 33324^T^, *C. eutactus* ATCC 27759^T^ and *L. multipara* DSM 3073^T^ were from NCBI.

| Strain | Accession number | Genome size (×10<sup>6</sup> bp) | G+C content (mol%) |
| --- | --- | --- | --- |
| Strain TF01-11<sup>T</sup> | LLKB00000000 | 3.61 | 36.79 |
| *R. intestinalis* DSM 14610<sup>T</sup> | ABYJ00000000 | 4.38 | 42.6 |
| *A. ethanolgignens* ATCC 33324<sup>T</sup> | LNAM00000000 | 3.66 | 41.0 |
| *C. eutactus* ATCC 27759<sup>T</sup> | ABEY00000000 | 3.10 | 43.1 |
| *L. multipara* DSM 3073<sup>T</sup> | AUJG00000000 | 2.87 | 35.3 |

**Table 3.**
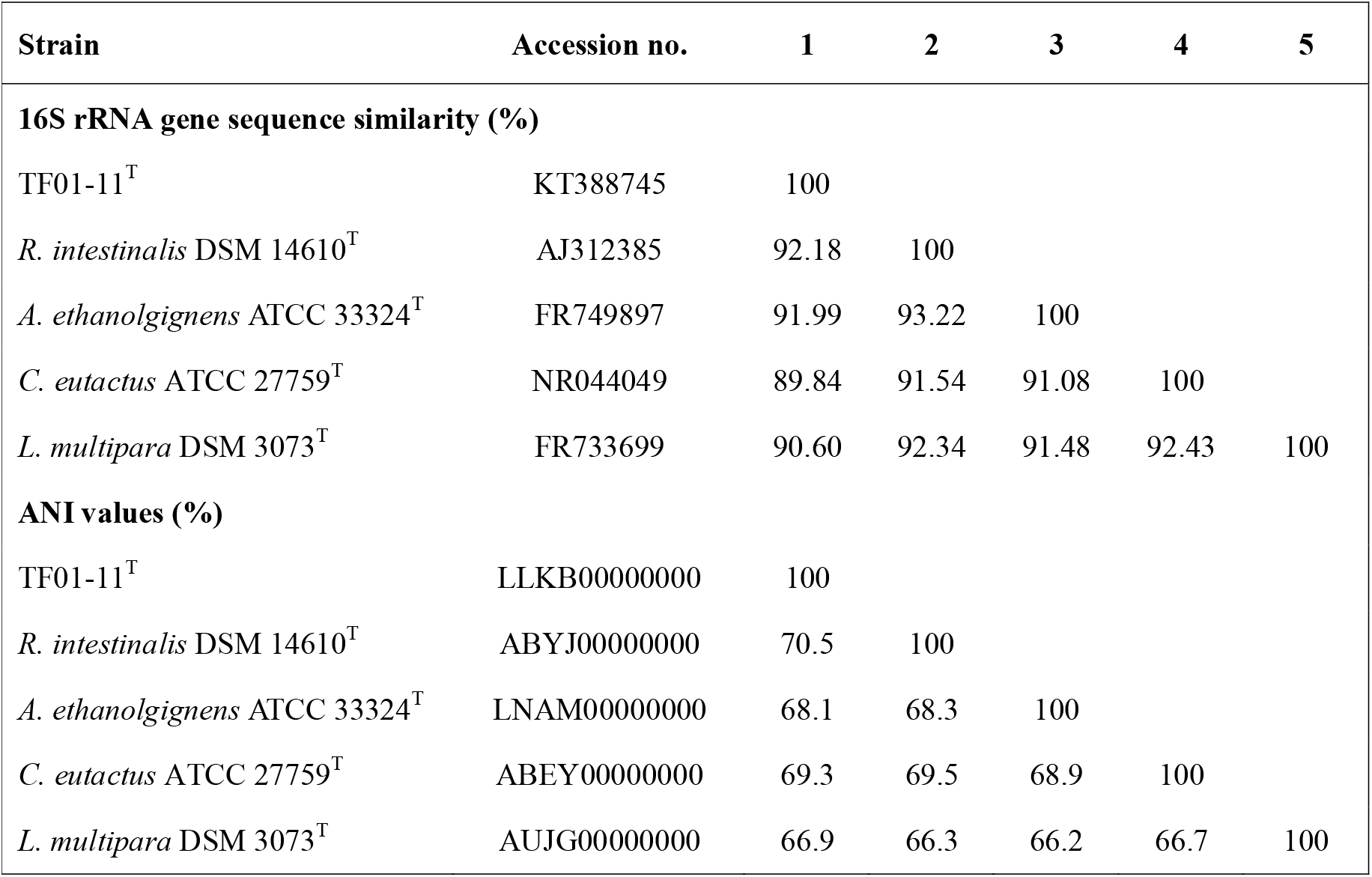
Levels of 16S rRNA gene sequence similarity and ANI values (in percentages) based on BLAST for strains TF01-11T and the most closely related members of the family *Lachnospiraceae.* Strains: TF01-11^T^; 2, *R. intestinalis* DSM 14610^T^; 3, *A. ethanolgignens* ATCC 33324^T^; 4, *C. eutactus* ATCC 27759^T^; 5, *L. multipara* DSM 3073^T^.

For RAST annotation with genome of strain TF01-11^T^, there were 11 genes associated with diaminopimelic acid synthesis, 7 genes associated with metabolism of polyamines, 12 genes associated with teichoic and lipoteichoic acids and 20 genes associated with metabolism of polar lipids, respectively, present in the genome (Table 4 and Table S2). No predicted gene sequences with recognizable similarity to those responsible for respiratory lipoquinones, mycolic acids or lipopolysaccharides.

**Table 4.**
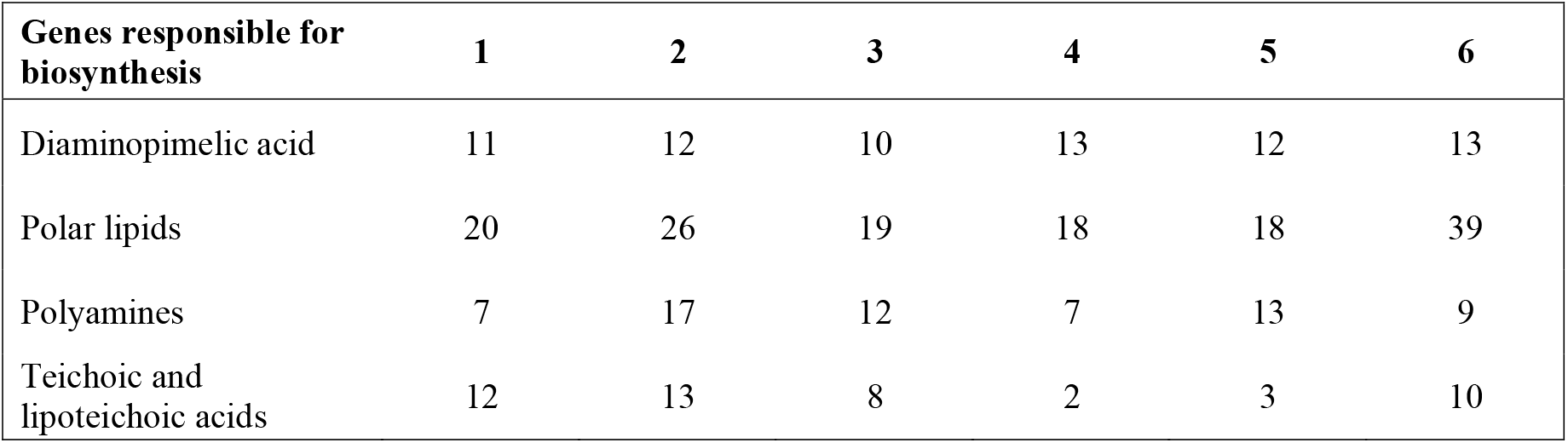
Number of genes identified in biosynthetic pathway from whole genome sequences of strain TF01-11^T^ and related organisms identified by RAST. Taxa: 1, TF01-11^T^; 2, *R. intestinalis* DSM 14610^T^; 3, *A. ethanolgignens* ATCC 33324^T^; 4, *C. eutactus* ATCC 27759^T^; 5, *L. multipara* DSM 3073^T^; 6, *Anaerostipes caccae* DSM 14662^T^. Data are for type strains. Numbers of genes identified for benzoquinones (ubiquinones, rhodoquinones, plastoquinones), naphthoquinones (menaquinones, demethylmenaquinones, monomethylmenaquinones, menathioquinones), lipopolysaccharides and mycolic acids were zero for all taxa studied.

| **Genes responsible for biosynthesis** | **1** | **2** | **3** | **4** | **5** | **6** |
| --- | --- | --- | --- | --- | --- | --- |
| Diaminopimelic acid | 11 | 12 | 10 | 13 | 12 | 13 |
| Polar lipids | 20 | 26 | 19 | 18 | 18 | 39 |
| Polyamines | 7 | 17 | 12 | 7 | 13 | 9 |
| Teichoic and lipoteichoic acids | 12 | 13 | 8 | 2 | 3 | 10 |

### Phenotypic and chemotaxonomic characteristics

Strain TF01-11^T^ was non-motile, Gram-stain-positive and non-spore-forming. No growth occurred under aerobic conditions. Cells were approximately 0.5–1.0 μm in width and 2.0–8.0 μm in length and occurring singly or in chains. Colonies on PYG agar plates were approximately 2.0 mm in diameter, grayish white, opaque, flat, smooth, dull and irregular with rhizoid margins after 48 h incubation at 37 °C under anaerobic conditions. The temperature for growth range from 30 °C to 42 °C (optimum 37 °C). The pH growth range was pH 6.0–8.5 (optimum pH 7.0). The NaCl tolerance range was 0 %–2.0 % (w/v). The major SCFAs produced in PYG broth were butyric and acetic acids. The main physiological and biochemical properties of strain TF01-11^T^ are given in Table 1 in comparison with closely related genera within family *Lachnospiraceae*. The predominant cellular fatty acid (>10 %) of strain TF01-11^T^ were C_16:0_ (37.9 %), C_14:0_ (16.4 %), C_13:0_ OH/iso-C_15:1_ H (11.1 %) and C_18:1_ *ω*9*c* (10.6 %) (Table S1, available in the online Supplementary Material), The whole-cell hydrolysate of strain TF01-11^T^ contained meso-diaminopimelic acid (m-DAP).

### Taxonomic conclusions

Base on genomic, phylogeny, phenotypic and chemotaxonomic characteristics of strain TF01-11^T^ presented above, we propose that this strain isolated from the faeces represents a novel species of a new genus distinct from other currently known species of family *Lachnospiraceae*, for which the name *Butyribacter intestini* gen. nov., sp. nov. is proposed. In addition, we propose reclassifying *Acetivibrio ethanolgignens* ATCC 33324^T^ within a new genus in the family *Lachnospiraceae*, *Acetanaerobacter ethanolgignens* gen. nov., comb. nov.

### Description of *Butyribacter* gen. nov.

*Butyribacter* (Bu.ty.ri.bac′ter. N.L. n. *acidum butyricum* butyric acid; N.L masc. n. *bacter* a rod; N.L. masc. n. *Butyribacter* a butyric acid-producing rod).

Gram-stain-negative, non-motile, non-spore-forming rods, about 2.0–8.0 μm long and 0.5–1.0 μm wide, occurring singly or in chains. Obligately anaerobic. Optimum growth temperature is approximately 37 °C. Butyric and acetic acids are the major metabolic end products in PYG broth. Meso-diaminopimelic acid is present in the hydrolysate of the peptidoglycan. The main fatty acids are C_16:0_, C_14:0_, C_13:0_ OH/iso-C_15:1_ H and C_18:1_ *ω*9*c*. The genome size is circa 3.6 Mbp. The genus is affiliated to the family *Lachnospiraceae*. The type species is *Butyribacter intestini*.

### Description of *Butyribacter intestini* sp. nov.

*Butyribacter intestini* (in.tes′ti.ni. L. gen. n. *intestini* of the gut, referring to the ecosystem of origin of the bacterium).

Cell morphology is the same as described for the genus. Colonies are approximately 2.0 mm in diameter, grayish white, opaque, flat, smooth, dull and irregular with rhizoid margins after 48 h at 37 °C. Growth occurs between 30 and 42 °C (optimum 37 °C) and at pH 6.0–8.5 (optimum pH 7.0–7.5). Positive result in tests for acid phosphatase (weak reaction) and *α*-glucosidase activities, but negative for alkaline phosphatase, esterase (C4), esterase lipase (C8), lipase (C14), leucine arylamidase, valine arylamidase, cystine arylamidase, trypsin, *α*-chymotrypsin, naphthol-AS-BI-phosphohydrolase, *α*-galactosidase, *β*-galactosidase, *β*-glucuronidase, *β*-glucosidase, *N*-acetyl-*β*-glucosaminidase, *α*-mannosidase and *β*-fucosidase. Acid is produced from glucose, ribose (week positive), galactose, fructose, methyl-D-glucopyranoside, cellobiose, maltose, melibiose, sucrose, starch, turanose, but not from xylose, adonitol, salicin, methyl-*β*-D-xylopyranoside, arabinose, glycerol, sorbose, dulcitol, melezitose, inositol, raffinose, mannitol, sorbitol, rhamnose, methyl-*α*-D-mannopyranoside, *N*-acetyl-glucosamine, amygdalin, arbutin, trehalose, inulin, glycogen, xylitol, gentiobiose, lyxose, tagatose, fucose, arabitol, gluconate, 2-ketogluconate and 5-ketogluconate. Indole is not produced. Gelatin is liquefied. Aesculin is not hydrolysed. There were 11 genes/proteins responsible for biosynthesis of DAP, including 4-hydroxy-tetrahydrodipicolinate reductase (EC 1.17.1.8) (1 gene), 4-hydroxy-tetrahydrodipicolinate synthase (EC 4.3.3.7) (1 gene), aspartate-semialdehyde dehydrogenase (EC 1.2.1.11) (1 gene), aspartokinase (EC 2.7.2.4) (1 gene), diaminopimelate decarboxylase (EC 4.1.1.20) (1 gene), diaminopimelate epimerase (EC 5.1.1.7) (1 gene), L,L-diaminopimelate aminotransferase (EC 2.6.1.83) (1 gene), *N*-acetyl-L, L-diaminopimelate deacetylase (EC 3.5.1.47) (2 genes), UDP-*N*-acetylmuramoylalanyl-D-glutamate--2,6-diaminopimelate ligase (EC 6.3.2.13) (1 gene), and UDP-*N*-acetylmuramoylalanyl-D-glutamyl-2,6-diaminopimelate--D-alanyl-D-alanine ligase (EC 6.3.2.10) (1 gene), 7 genes/proteins responsible for biosynthesis of polyamines, including 5’-methylthioadenosine nucleosidase (EC 3.2.2.16) @ *S*-adenosylhomocysteine nucleosidase (EC 3.2.2.9) (1 gene), ABC transporter, periplasmic spermidine putrescine-binding protein PotD (TC 3.A.1.11.1) (1 gene), arginine decarboxylase (EC 4.1.1.19) / lysine decarboxylase (EC 4.1.1.18) (1 gene), carbamate kinase (EC 2.7.2.2) (1 gene), putrescine transport ATP-binding protein PotA (TC 3.A.1.11.1) (1 gene), spermidine Putrescine ABC transporter permease component PotB (TC 3.A.1.11.1) (1 gene), and spermidine putrescine ABC transporter permease component potC (TC..3.A.1.11.1) (1 gene), 12 genes/protein responsible for biosynthesis of teichoic and lipoteichoic acids, including 2-*C*-methyl-D-erythritol 4-phosphate cytidylyltransferase (EC 2.7.7.60) (2 genes), CDP-glycerol:poly(glycerophosphate) glycerophosphotransferase (EC 2.7.8.12) (2 genes), membrane protein involved in the export of O-antigen, teichoic acid lipoteichoic acids (1 gene), minor teichoic acid biosynthesis protein GgaB (1 gene), *N*-acetylmannosaminyltransferase (EC 2.4.1.187) (1 gene), teichoic acid export ATP-binding protein TagH (EC 3.6.3.40) (3 genes), teichoic acid glycosylation protein (1 gene), and teichoic acid translocation permease protein TagG (1 gene), and 20 genes/protein responsible for biosynthesis of polar lipids, including 1-acyl-sn-glycerol-3-phosphate acyltransferase (EC 2.3.1.51) (2 genes), acyl carrier protein (2 genes), acyl-phosphate:glycerol-3-phosphate O-acyltransferase PlsY (1 gene), alcohol dehydrogenase (EC 1.1.1.1) (1 gene), acetaldehyde dehydrogenase (EC 1.2.1.10) (1 gene), aldehyde dehydrogenase (EC 1.2.1.3) (1 gene), cardiolipin synthetase (EC 2.7.8.-) (1 gene), CDP-diacylglycerol--glycerol-3-phosphate 3-phosphatidyltransferase (EC 2.7.8.5) (1 gene), CDP-diacylglycerol--serine O-phosphatidyltransferase (EC 2.7.8.8) (1 gene), dihydroxyacetone kinase family protein (1 gene), glycerate kinase (EC 2.7.1.31) (1 gene), glycerol-3-phosphate dehydrogenase (EC 1.1.5.3) (1 gene), glycerol-3-phosphate dehydrogenase [NAD(P)^+^] (EC 1.1.1.94) (1 gene), phosphate:acyl-ACP acyltransferase PlsX (1 gene), phosphatidate cytidylyltransferase (EC 2.7.7.41) (1 gene), phosphatidylglycerophosphatase B (EC 3.1.3.27) (1 gene), and phosphatidylserine decarboxylase (EC 4.1.1.65) (1 gene). There are no genes responsible for biosynthesis of respiratory lipoquinones, mycolic acids or lipopolysaccharides.

The type strain is TF01-11^T^ (CGMCC 1.5203^T^ **=** CGMCC 10984^T^ = DSM 105140^T^), isolated from the faeces obtained from a 17-year-old Chinese female. The DNA G+C content of the type strain is 36.79 mol%.

### Description of *Acetanaerobacter* gen. nov.

*Acetanaerobacter* (A.cet.an.ae.ro. bac′ter. L. n. *acetum* vinegar; Gr. pref. *an* not; Gr. masc. n. *aer* air; N.L masc. n. *bacter* a rod; N.L. masc. n. *Acetanaerobacter* vinegar-producing anaerobic rod).

Cells are nonsporeforming, motile, Gram-stain-negative rods, obligately anaerobic, which do not grow under microaerophilic or aerobic conditions. Growth occurs between 30 and 42 °C (optimum 37 °C) and at pH 7.0–9.0 (optimum pH 7.0). Acetic and lactate acids are the major metabolic end products in PYG broth. Major fatty acids are C_14:0_, C_16:0_ and C_18:1_ *ω*9*c*. The genome size is circa 3.7 Mbp. The genomic DNA G+C content of the type species is 41.0 mol%. The type species is *Acetanaerobacter ethanolgignens.*

### Description of *Acetanaerobacter ethanolgignens* comb. nov.

The description of the species is given by Robinson & Ritchie (1881) and determined from this study. Cells are about 0.5–0.9 μm long and 1.5–2.5 μm wide, occurring singly, in pairs, and often in short chains. Colonies are yellowish-white, circular, convex, smooth, translucent, and 0.5 to 1.5 mm in diameter on PYG plate after 2–3 days of cultivation. The fermentation products is acetic acid, ethanol, hydrogen, and carbon dioxide. Positive result in tests for leucine arylamidase activity, but negative for acid phosphatise, *α*-glucosidase activities, alkaline phosphatase, esterase (C4), esterase lipase (C8), lipase (C14), valine arylamidase, cystine arylamidase, trypsin, *α*-chymotrypsin, naphthol-AS-BI-phosphohydrolase, *α*-galactosidase, *β*-galactosidase, *β*-glucuronidase, *β*-glucosidase, *N*-acetyl-*β*-glucosaminidase, *α*-mannosidase and *β*-fucosidase. Acid is produced from glucose, mannose, lactose, maltose, salicin, mannose, cellobiose (weak positive), raffinose (weak positive), fructose, galactose, mannitol, pyruvate and starch (weak positive), but not from xylose, arabinose, glycerol, melezitose, sorbitol, rhamnose, trehalose, adonitol, lactate, raffinose, ribose and sucrose. Indole is not produced. Gelatin is liquefied. Aesculin is not hydrolysed.

The type strain is 77-6^T^ (= ATCC 33324^T^ = DSM 3005^T^)

## Figure legends

**Fig. 1.** Neighbour-joining phylogenetic tree based on 16S rRNA gene sequence of strain TF01-11^T^ and related type species of the family *Lachnospiraceae* and *Ruminococcaceae*. Bootstrap values (> 90 %) based on 1000 replications are shown at branch nodes. *Clostridium butyricum* is used as an out-group. Bar, 20 % nucleotide sequence divergence.

## Acknowledgements

This work was supported by grants from National Key Research and Development Program of China (No. 2018YFC1313800) and Natural Science Foundation of Guangdong Province, China (No. 2019B020230001). We are grateful to Professor Fang Chengxiang for his support in technical advice. We also thank the colleagues at BGI-Shenzhen for sample collection, and discussions, and China National Genebank (CNGB) Shenzhen for DNA extraction, library construction, sequencing.

## Notes

### Competing Interest Statement

The authors have declared no competing interest.

