## Supplementary Data for "*Butyribacter intestini* gen. nov., sp. nov., a butyric acid-producing bacterium of the family *Lachnospiraceae* isolated from the human faeces, and reclassification of *Acetivibrio ethanolgignens* as *Acetanaerobacter ethanolgignens* gen. nov., comb. nov"

**Table S1.** Cellular fatty acid composition of strain TF01-11$^T$ and closely related species. Strains: 1, TF01-11$^T$; 2, *R. intestinalis* DSM 14610$^T$; 3, *A. ethanolgignens* ATCC 33324$^T$; 4, *L. multipara* DSM 3073$^T$; Data were obtained in this study. –, Not detected.

| **Fatty acids** | **1** | **2** | **3** | **4** |
|---|---|---|---|---|
| $C_{10:0}$ | – | – | 1.1 | – |
| $C_{12:0}$ | – | 1.4 | 1.7 | – |
| $C_{13:1}$ | 5.1 | – | 1.1 | 1.4 |
| $C_{14:0}$ | **16.4** | 9.2 | **13.3** | 8.0 |
| $C_{13:0}$ 3OH/iso-$C_{15:1}$ H | **11.1** | – | 2.3 | 3.8 |
| $C_{16:1}$ $\omega 9c$ | – | – | 3.8 | – |
| $C_{16:1}$ $\omega 7c$/ $C_{16:1}$ $\omega 6c$ | – | 6.1 | 5.0 | – |
| $C_{16:0}$ | **37.9** | **33.9** | **32.5** | **44.0** |
| $C_{17:1}$ $\omega 8c$ | 1.31 | **–** | **–** | **–** |
| Iso-$C_{17:1}$ I/anteiso B | – | 1.2 | 3.9 | 7.9 |
| $C_{18:1}$ $\omega 9c$ | **10.6** | **19.9** | **18.2** | **16.0** |
| $C_{18:1}$ $\omega 7c$ | 1.9 | 8.2 | 5.6 | 3.3 |
| $C_{18:1}$ $\omega 6c$ | – | **13.8** | 2.8 | – |
| $C_{18:0}$ | 2.4 | 4.8 | 5.2 | 5.2 |
| $C_{18:0}$ 2 OH | – | – | – | 2.3 |
| Iso-$C_{19:0}$ | 4.3 | – | – | |
| Iso-$C_{19:1}$ I | – | – | – | 1.7 |
| $C_{19:1}$ $\omega 9c$/$C_{19:1}\omega 11c$ | 3.4 | – | – | – |
| Anteiso-$C_{19:0}$ | – | – | – | 2.0 |

**Table S2.** Levels of 16S rRNA gene sequence similarity and ANI values (in percentages) based on BLAST for strains TF01-11$^{T}$ and the most closely related members of the family *Lachnospiraceae*. ND, not data.

Strains: TF01-11$^{T}$; 2, *R. intestinalis* DSM 14610$^{T}$; 3, *A. ethanolgignens* ATCC 33324$^{T}$; 4, *C. eutactus* ATCC 27759$^{T}$; 5, *L. multipara* DSM 3073$^{T}$;

| **Strain** | **Accession no.** | **1** | **2** | **3** | **4** | **5** |
|---|---|---|---|---|---|---|
| **16S rRNA gene sequence similarity (%)** | | | | | | |
| TF01-11$^{T}$ | KT388745 | 100 | | | | |
| *R. intestinalis* DSM 14610$^{T}$ | AJ312385 | 92.18 | 100 | | | |
| *A. ethanolgignens* ATCC 33324$^{T}$ | FR749897 | 91.99 | 93.22 | 100 | | |
| *C. eutactus* ATCC 27759$^{T}$ | NR044049 | 89.84 | 91.54 | 91.08 | 100 | |
| *L. multipara* DSM 3073$^{T}$ | FR733699 | 90.60 | 92.34 | 91.48 | 92.43 | 100 |
| **ANI values (%)** | | | | | | |
| TF01-11$^{T}$ | LLKB00000000 | 100 | | | | |
| *R. intestinalis* DSM 14610$^{T}$ | ABYJ00000000 | 70.5 | 100 | | | |
| *A. ethanolgignens* ATCC 33324$^{T}$ | LNAM00000000 | 68.1 | 68.3 | 100 | | |
| *C. eutactus* ATCC 27759$^{T}$ | ABEY00000000 | 69.3 | 69.5 | 68.9 | 100 | |
| *L. multipara* DSM 3073$^{T}$ | AUJG00000000 | 66.9 | 66.3 | 66.2 | 66.7 | 100 |

**Table S3.** The specific genes/protein related to biosynthesis of DAP, polar lipids, polyamines and lipoteichoic and teichoic acids and their positions in the genome in comparation of strain TF01-11$^{T}$ and related organisms identified by Rapid Annotation Subsystem Technology (RAST).

| **Gene /Protein related to DAP** | **Strain TF01-11$^{T}$** | **ATCC 33324$^{T}$** | **DSM 14610$^{T}$** | **ATCC 27759$^{T}$** | **DSM 3073$^{T}$** | **DSM 14662$^{T}$** |
|---|---|---|---|---|---|---|
| 2,3,4,5-tetrahydropyridine-2,6-dicarboxylate N-acetyltransferase (EC 2.3.1.89) | – | LNAM01000219: 69918..70397 | – | – | – | – |
| 4-hydroxy-tetrahydrodipicolinate reductase (EC 1.17.1.8) | LLKB01000001.1: 891848..892603 | LNAM01000037: 21206..21961 | ABYJ02000093.1: 29320..30147 | ABEY02000020.1: 68740..69567 | AUJG01000001.1: 207142..207897 | ABAX03000001.1: 64350..63595 |
| 4-hydroxy-tetrahydrodipicolinate synthase (EC 4.3.3.7) | LLKB01000001.1: 890818..891711 | LNAM01000037: 20286..21170 | ABYJ02000093.1: 28385..29275 | ABEY02000020.1: 67706..68590 | AUJG01000001.1: 206223..207107 | ABAX03000014.1: 342777..340588<br>ABAX03000001.1: 65247..64363 |
| Acetylornithine aminotransferase (EC 2.6.1.11) / N-succinyl-L,L-diaminopimelate aminotransferase (EC 2.6.1.17) | – | – | – | – | AUJG01000005.1: 63513..62329 | – |
| Aspartate-semialdehyde dehydrogenase (EC 1.2.1.11) | LLKB01000005.1: 266333..265236 | LNAM01000001: 11487..10399 | ABYJ02000204.1: 3750..4841 | ABEY02000032.1: 231619..232710 | AUJG01000002.1: 64603..65691 | ABAX03000012.1: 122581..123666 |
| Aspartokinase (EC 2.7.2.4) | LLKB01000001.1: 615848..617167 | LNAM01000001: 7468..6257 | ABYJ02000070.1: 8837..7518 | ABEY02000027.1: 73295..74656<br>ABEY02000025.1: 240234..241439 | AUJG01000020.1: 17450..16131 | ABAX03000024.1: 415789..416991<br>ABAX03000012.1: 235578..236048 |
| Diaminopimelate decarboxylase (EC 4.1.1.20) | LLKB01000006.1: 118004..119272 | LNAM01000057: 11038..12303 | ABYJ02000062.1: 2599..2159<br>ABYJ02000062.1: 2157..1315 | ABEY02000007.1: 97145..95889 | AUJG01000015.1: 62520..61300<br>AUJG01000002.1: 362371..361094 | ABAX03000013.1: 59804..58554 |
| Diaminopimelate epimerase (EC 5.1.1.7) | LLKB01000005.1: 31112..30276 | LNAM01000164: 81730..82569 | ABYJ02000147.1: 7714..6848<br>ABYJ02000047.1: 24811..23966 | ABEY02000022.1: 204226..203357 | AUJG01000014.1: 69657..68785 | ABAX03000012.1: 736168..735332 |
| L,L-diaminopimelate aminotransferase (EC 2.6.1.83) | LLKB01000001.1: 1371235..1370015 | LNAM01000079: 41120..39906 | ABYJ02000145.1: 8628..7414 | ABEY02000022.1: 203355..202138 | AUJG01000005.1: 24511..23297 | ABAX03000001.1: 19692..18478 |
| Meso-diaminopimelate D-dehydrogenase (EC 1.4.1.16) | – | – | – | – | AUJG01000010.1: 84685..83699 | ABAX03000021.1: 2432..3418 |
| N-acetyl-L,L-diaminopimelate deacetylase (EC 3.5.1.47) | LLKB01000001.1: 950282..951439<br>LLKB01000005.1: 14428..13250 | – | ABYJ02000174.1: 11764..10631 | ABEY02000001.1: 115756..114545<br>ABEY02000025.1: 103606..102083<br>ABEY02000018.1: 38609..36987 | – | ABAX03000010.1: 47209..46070 |
| UDP-N-acetylmuramoylalanyl-D-glutamate--2,6-diaminopimelate ligase (EC 6.3.2.13) | LLKB01000005.1: 30186..28717 | LNAM01000164: 82583..84040 | ABYJ02000250.1: 1295..993<br>ABYJ02000147.1: 6606..5086 | ABEY02000006.1: 44415..43648 | AUJG01000018.1: 32698..31196 | ABAX03000012.1: 735307..733847 |
| UDP-N-acetylmuramoylalanyl-D-glutamyl-2,6-diaminopimelate--D-alanyl-D-alanine ligase (EC 6.3.2.10) | LLKB01000005.1: 280599..279211 | LNAM01000164: 84043..85431 | – | ABEY02000022.1: 88858..87371 | AUJG01000005.1: 56828..55449 | ABAX03000005.1: 6685..8040 |

| Gene /Protein related to polar lipids | Strain TF01-11$^T$ | ATCC 33324$^T$ | DSM 14610$^T$ | ATCC 27759$^T$ | DSM 3073$^T$ | DSM 14662$^T$ |
|---|---|---|---|---|---|---|
| 1-acyl-sn-glycerol-3-phosphate acyltransferase (EC 2.3.1.51) | LLKB01000005.1: 94808..94071 | LNAM01000101: 54051..53296 | ABYJ02000168.1: 15753..14998 | ABEY02000006.1: 379016..378336 | AUJG01000003.1: 114048..114800 | ABAX03000037.1: 64326..63637 |
| | LLKB01000005.1: 1459117..1458398 | | ABYJ02000055.1: 46968..46234 | ABEY02000006.1: 592105..592845 | | ABAX03000024.1: 318948..318211 |
| Acyl carrier protein | LLKB01000005.1: 134289..134071 | LNAM01000046: 24658..24894 | ABYJ02000188.1: 12731..12501 | ABEY02000018.1: 102596..102844 | AUJG01000010.1: 23499..23735 | ABAX03000024.1: 347378..347611 |
| | LLKB01000006.1: 75421..75648 | LNAM01000160: 23002..23250 | ABYJ02000005.1: 19375..19145 | ABEY02000006.1: 394572..394339 | AUJG01000002.1: 264720..264950 | |
| | | LNAM01000175: 87574..87353 | | | | |
| Acyl-phosphate:glycerol-3-phosphate O-acyltransferase PlsY | LLKB01000001.1: 1485909..1485277 | LNAM01000068: 52532..51900 | ABYJ02000238.1: 5911..5258 | ABEY02000002.1: 35573..34914 | AUJG01000002.1: 290459..291124 | ABAX03000012.1: 746057..745431 |
| | | | ABYJ02000056.1: 4435..3815 | | | |
| Alcohol dehydrogenase (EC 1.1.1.1) | LLKB01000001.1: 1020838..1019750 | – | – | ABEY02000002.1: 33320..22617 | AUJG01000002.1: 114064..115251 | ABAX03000039.1: 127478..126333 |
| | | | | | | ABAX03000037.1: 104962..103910 |
| | | | | | | ABAX03000037.1: 133315..132260 |
| | | | | | | ABAX03000024.1: 138615..137569 |
| | | | | | | ABAX03000014.1: 99513..98503 |
| | | | | | | ABAX03000014.1: 104291..103119 |
| | | | | | | ABAX03000014.1: 345082..343865 |
| | | | | | | ABAX03000013.1: 7469..6459 |
| | | | | | | ABAX03000012.1: 18810..19961 |
| | | | | | | \|ABAX03000012.1: 22950..24134 |
| | | | | | | ABAX03000012.1: 247376..248428 |
| | | | | | | ABAX03000012.1: 453543..452503 |
| | | | | | | ABAX03000001.1: 176544..175456 |
| Alcohol dehydrogenase (EC 1.1.1.1); Acetaldehyde dehydrogenase (EC 1.2.1.10) | LLKB01000005.1: 1128303..1127092 | LNAM01000090: 10449..9271 | ABYJ02000187.1: 4980..6197 | ABEY02000025.1: 175328..176545 | AUJG01000004.1: 69398..68190 | ABAX03000012.1: 688831..687656 |
| | | LNAM01000208: 45801..43180 | | | AUJG01000004.1: 118447..121083 | |
| Aldehyde dehydrogenase (EC 1.2.1.3) | LLKB01000001.1: 82173..80797 | LNAM01000199: 3302..1923 | – | – | – | ABAX03000012.1: 301799..303238 |
| | | | | | | ABAX03000012.1: 381267..379897 |
| Cardiolipin synthetase (EC 2.7.8.-) | LLKB01000001.1: 1466186..1465314 | LNAM01000113: 15358..13697 | ABYJ02000181.1: 897..706 | ABEY02000022.1: 48213..46684 | AUJG01000011.1: 42121..43680 | ABAX03000038.1: 102403..103965 |
| | LLKB01000005.1: 63476..61824 | | ABYJ02000181.1: 9675..8461 | | | |

| | | | | | | |
|---|---|---|---|---|---|---|
| | | | ABYJ02000181.1: 18047..17064<br>ABYJ02000158.1: 17566..15995<br>ABYJ02000152.1: 58128..56593 | | | |
| CDP-diacylglycerol--glycerol-3-phosphate 3-phosphatidyltransferase (EC 2.7.8.5) | LLKB01000005.1: 141182..140634 | LNAM01000046: 14967..15503 | ABYJ02000181.1: 2024..1464<br>ABYJ02000044.1: 58194..58736 | ABEY02000003.1: 130088..129564 | AUJG01000002.1: 215432..215974 | ABAX03000038.1: 164188..164682<br>ABAX03000012.1: 320025..319465 |
| CDP-diacylglycerol--serine O-phosphatidyltransferase (EC 2.7.8.8) | LLKB01000006.1: 196056..196775 | – | – | ABEY02000003.1: 105452..104745 | – | ABAX03000016.1: 27420..26809 |
| Diacylglycerol kinase (EC 2.7.1.107) | – | LNAM01000057: 45914..45540 | – | – | – | – |
| Dihydroxyacetone kinase family protein | LLKB01000005.1: 394221..392560 | LNAM01000113: 31994..30324 | ABYJ02000108.1: 6563..6312<br>ABYJ02000044.1: 29676..31373 | ABEY02000003.1: 161296..159605 | AUJG01000011.1: 89949..90275<br>AUJG01000002.1: 356764..355052 | ABAX03000002.1: 122226..120586 |
| Glycerate kinase (EC 2.7.1.31) | LLKB01000005.1: 1389832..1388693 | LNAM01000005: 2379..3518 | ABYJ02000081.1: 6301..7422 | ABEY02000029.1: 52091..50805 | AUJG01000002.1: 50028..48886 | ABAX03000038.1: 145464..144349<br>ABAX03000037.1: 35793..34639<br>ABAX03000037.1: 42966..42400<br>ABAX03000010.1: 196597..195461 |
| Glycerol kinase (EC 2.7.1.30) | – | LNAM01000176: 4341..5837 | ABYJ02000094.1: 38045..36549 | ABEY02000006.1: 39199..37712 | AUJG01000014.1: 59383..60888 | ABAX03000013.1: 25425..23929 |
| Glycerol-1-phosphate dehydrogenase [NAD(P)] (EC 1.1.1.261) | – | – | – | – | – | ABAX03000024.1: 25782..27074<br>ABAX03000012.1: 455990..454701<br>ABAX03000003.1: 65986..64703 |
| Glycerol-3-phosphate dehydrogenase (EC 1.1.5.3) | LLKB01000001.1: 1153693..1153361 | LNAM01000176: 5957..7393<br>LNAM01000202: 4102..5820 | ABYJ02000094.1: 35581..34082 | – | – | ABAX03000021.1: 11232..12671 |
| Glycerol-3-phosphate dehydrogenase [NAD(P)+] (EC 1.1.1.94) | LLKB01000001.1: 1485198..1484179 | LNAM01000068: 51882..50869 | ABYJ02000238.1: 5253..4843<br>ABYJ02000238.1: 4841..4239<br>ABYJ02000054.1: 12162..14855 | ABEY02000002.1: 34435..33446 | AUJG01000002.1: 291117..292127 | ABAX03000021.1: 14307..15350<br>ABAX03000012.1: 745421..744405 |
| Phosphate:acyl-ACP acyltransferase PlsX | LLKB01000005.1: 135308..134301 | LNAM01000046: 23632..24642 | ABYJ02000188.1: 13776..12736 | ABEY02000003.1: 20139..21149 | AUJG01000002.1: 263634..264647 | ABAX03000002.1: 99849..98839 |
| Phosphatidate cytidylyltransferase (EC 2.7.7.41) | LLKB01000001.1: 929173..929964 | LNAM01000001: 84925..85731 | ABYJ02000224.1: 3587..4393 | ABEY02000018.1: 15638..14841 | AUJG01000007.1: 9819..9010 | ABAX03000012.1: 371060..370245 |
| Phosphatidylglycerophosphatase B (EC 3.1.3.27) | LLKB01000005.1: 463804..463277 | – | ABYJ02000173.1: 6108..5563<br>ABYJ02000038.1: 47582..47097 | ABEY02000027.1: 30004..30558 | AUJG01000005.1: 41676..41044<br>AUJG01000002.1: 308417..308758 | – |
| Phosphatidylserine decarboxylase (EC 4.1.1.65) | LLKB01000006.1: 196850..197710 | – | – | ABEY02000003.1: 104666..103677 | – | ABAX03000016.1: 26796..25915 |

| **Gene /Protein related to polyamine** | **Strain TF01-11$^T$** | **ATCC 33324$^T$** | **DSM 14610$^T$** | **ATCC 27759$^T$** | **DSM 3073$^T$** | **DSM 14662$^T$** |
|---|---|---|---|---|---|---|
| 5'-methylthioadenosine nucleosidase (EC 3.2.2.16) @ S-adenosylhomocysteine | LLKB01000005.1: 933531..932845 | – | ABYJ02000063.1: 4273..3620 | ABEY02000029.1: 19934..19308 | AUJG01000008.1: 56341..57033 | ABAX03000024.1: 36088..36804 |

| | | | | | | |
|---|---|---|---|---|---|---|
| nucleosidase (EC 3.2.2.9) | | | | ABEY02000029.1: 19934..19308 | AUJG01000008.1: 56341..57033 | ABAX03000012.1: 467287..467919 |
| ABC transporter, periplasmic spermidine putrescine-binding protein PotD (TC 3.A.1.11.1) | LLKB01000006.1: 171320..172402 | LNAM01000188: 11833..9944 | ABYJ02000114.1: 14238..13471<br>ABYJ02000114.1: 13467..13426<br>ABYJ02000114.1: 13422..13411<br>ABYJ02000114.1: 13411..12371 | ABEY02000001.1: 109509..107881 | – | – |
| Agmatine deiminase (EC 3.5.3.12) | – | – | ABYJ02000093.1: 38640..39863 | – | AUJG01000024.1: 6045..7178 | – |
| Arginine decarboxylase (EC 4.1.1.19) / Lysine decarboxylase (EC 4.1.1.18) | LLKB01000006.1: 197812..199482 | LNAM01000193: 11018..9597 | ABYJ02000093.1: 33558..35159<br>ABYJ02000067.1: 12063..13571 | ABEY02000007.1: 111988..110339 | AUJG01000024.1: 1010..2452<br>AUJG01000008.1: 2037..3563 | ABAX03000014.1: 403151..401772 |
| Arginine/ornithine antiporter ArcD | – | LNAM01000197: 35739..37151 | ABYJ02000122.1: 4433..3342 | – | AUJG01000001.1: 486447..487055<br>AUJG01000017.1: 30780..30178 | – |
| Carbamate kinase (EC 2.7.2.2) | LLKB01000001.1: 1175095..1176027 | LNAM01000164: 13022..12105<br>LNAM01000197: 34632..35546 | ABYJ02000162.1: 6928..5996 | – | AUJG01000018.1: 42064..41126 | ABAX03000039.1: 106418..107362 |
| Carboxynorspermidine decarboxylase, putative (EC 4.1.1.-) | – | – | ABYJ02000093.1: 37535..38668 | – | AUJG01000024.1: 4674..5903 | – |
| Carboxynorspermidine dehydrogenase, putative (EC 1.1.1.-) | | | ABYJ02000093.1: 36188..37447 | | AUJG01000024.1: 3343..4602 | |
| N-carbamoylputrescine amidase (3.5.1.53) | – | – | ABYJ02000093.1: 39941..40831 | – | AUJG01000024.1: 7935..8819<br>AUJG01000002.1: 7869..8666 | – |
| Putrescine transport ATP-binding protein PotA (TC 3.A.1.11.1) | LLKB01000006.1: 168610..169680 | LNAM01000153: 110671..109595<br>LNAM01000188: 14255..12675<br>LNAM01000219: 1483..3204 | ABYJ02000115.1: 1430..357 | ABEY02000001.1: 112736..111288 | – | ABAX03000038.1: 20351..21154<br>ABAX03000012.1: 442997..441312<br>ABAX03000004.1: 37815..36688 |
| Putrescine transport ATP-binding protein PotG (TC 3.A.1.11.2) | – | LNAM01000219: 31196..31972 | – | – | – | – |
| Spermidine Putrescine ABC transporter permease component PotB (TC 3.A.1.11.1) | LLKB01000006.1: 169696..170532 | LNAM01000188: 12675..11830 | ABYJ02000114.1: 14618..14232 | ABEY02000001.1: 111190..110390 | – | ABAX03000004.1: 36707..35856 |
| Spermidine Putrescine ABC transporter permease component potC (TC..3.A.1.11.1) | LLKB01000006.1: 170526..171323 | – | – | ABEY02000001.1: 110390..109596 | – | ABAX03000004.1: 35843..35043 |
| Spermidine synthase (EC 2.5.1.16) | – | LNAM01000156: 28318..29931 | ABYJ02000093.1: 35227..36090 | – | AUJG01000024.1: 2467..3324 | – |
| Transcriptional regulator, MerR family, near polyamine transporter | – | LNAM01000188: 14804..14271 | ABYJ02000115.1: 2039..1500 | – | – | – |

| **Gene /Protein related to Teichoic and lipoteichoic acids** | **Strain TF01-11$^T$** | **ATCC 33324$^T$** | **DSM 14610$^T$** | **ATCC 27759$^T$** | **DSM 3073$^T$** | **DSM 14662$^T$** |
|---|---|---|---|---|---|---|
| 1,2-diacylglycerol 3-glucosyltransferase (EC 2.4.1.157); diglucosyldiacylglycerol synthase (LTA membrane anchor synthesis); putative | – | — | ABYJ02000056.1: 5571..4432 | – | – | – |

| | | | | | | |
|---|---|---|---|---|---|---|
| 2-C-methyl-D-erythritol 4-phosphate cytidylyltransferase (EC 2.7.7.60) | LLKB01000005.1: 307612..306881<br>LLKB01000006.1: 156385..157089 | LNAM01000166: 23857..24561 | ABYJ02000192.1: 40539..41240<br>ABYJ02000069.1: 40611..39850 | – | – | ABAX03000027.1: 1659..2450 |
| CDP-glycerol:poly(glycerophosphate) glycerophosphotransferase (EC 2.7.8.12) | LLKB01000005.1: 475397..474219<br>LLKB01000006.1: 142480..143499 | LNAM01000079: 32976..31843<br>LNAM01000225: 5154..3973<br>LNAM01000079: 32976..31843 | – | – | – | ABAX03000001.1: 187884..186730<br>ABAX03000001.1: 193383..191245<br>ABAX03000001.1: 197038..195872 |
| CDP-glycerol: N-acetyl-beta-D-mannosaminyl-1,4-N-acetyl-D-glucosaminyldiphosphoundecaprenyl glycerophosphotransferase | – | – | ABYJ02000069.1: 43527..42250 | – | – | ABAX03000001.1: 201742..198386 |
| Glycosyltransferase LafA, responsible for the formation of Glc-DAG | – | LNAM01000155: 11554..12750 | – | – | – | – |
| Glycosyltransferase LafB, responsible for the formation of Gal-Glc-DAG | – | LNAM01000046: 7961..9022 | – | – | – | – |
| Membrane protein involved in the export of O-antigen, teichoic acid lipoteichoic acids | LLKB01000006.1: 256386..257852 | – | – | – | – | – |
| Minor teichoic acid biosynthesis protein GgaB | LLKB01000007.1: 66641..70336 | – | – | – | – | – |
| N-acetylmannosaminyltransferase (EC 2.4.1.187) | LLKB01000001.1: 113955..114749 | LNAM01000049: 17926..17171 | ABYJ02000039.1: 47789..47061<br>ABYJ02000003.1: 26093..25368 | – | – | – |
| Putative polyribitolphosphotransferase | – | – | ABYJ02000069.1: 46382..44628 | – | – | – |
| Teichoic acid biosynthesis protein | – | – | – | – | – | ABAX03000001.1: 195868..193412<br>ABAX03000001.1: 204601..207144<br>ABAX03000001.1: 207162..209489 |
| Teichoic acid export ATP-binding protein TagH (EC 3.6.3.40) | LLKB01000001.1: 1355716..1354976<br>LLKB01000005.1: 868414..867473<br>LLKB01000006.1: 240871..242067 | LNAM01000225: 7590..6847 | ABYJ02000069.1: 16969..15665<br>ABYJ02000069.1: 41459..40608<br>ABYJ02000069.1: 80501..79227 | ABEY02000032.1: 18825..19772<br>ABEY02000006.1: 446795..445569 | – | ABAX03000001.1: 202494..201751<br>ABAX03000001.1: 202494..201751 |
| Teichoic acid glycosylation protein | LLKB01000006.1: 262497..262925 | – | ABYJ02000039.1: 70515..70054 | – | AUJG01000001.1: 18248..17307 | – |
| Teichoic acid translocation permease protein TagG | LLKB01000006.1: 239972..240799 | – | ABYJ02000069.1: 17773..16988<br>ABYJ02000069.1: 81373..80540 | – | AUJG01000025.1: 15191..13956<br>AUJG01000030.1: 2582..2112 | – |

**Supplementary Fig. S1.** Maximum-likelihood phylogenetic tree based on 16S rRNA gene sequence of strain TF01-11$^{T}$ and related type species of the family *Lachnospiraceae* and *Ruminococcaceae*. Bootstrap values (>90 %) based on 1000 replications are shown at branch nodes. *Clostridium butyricum* is used as an out-group. Bar, 2 % nucleotide sequence divergence.

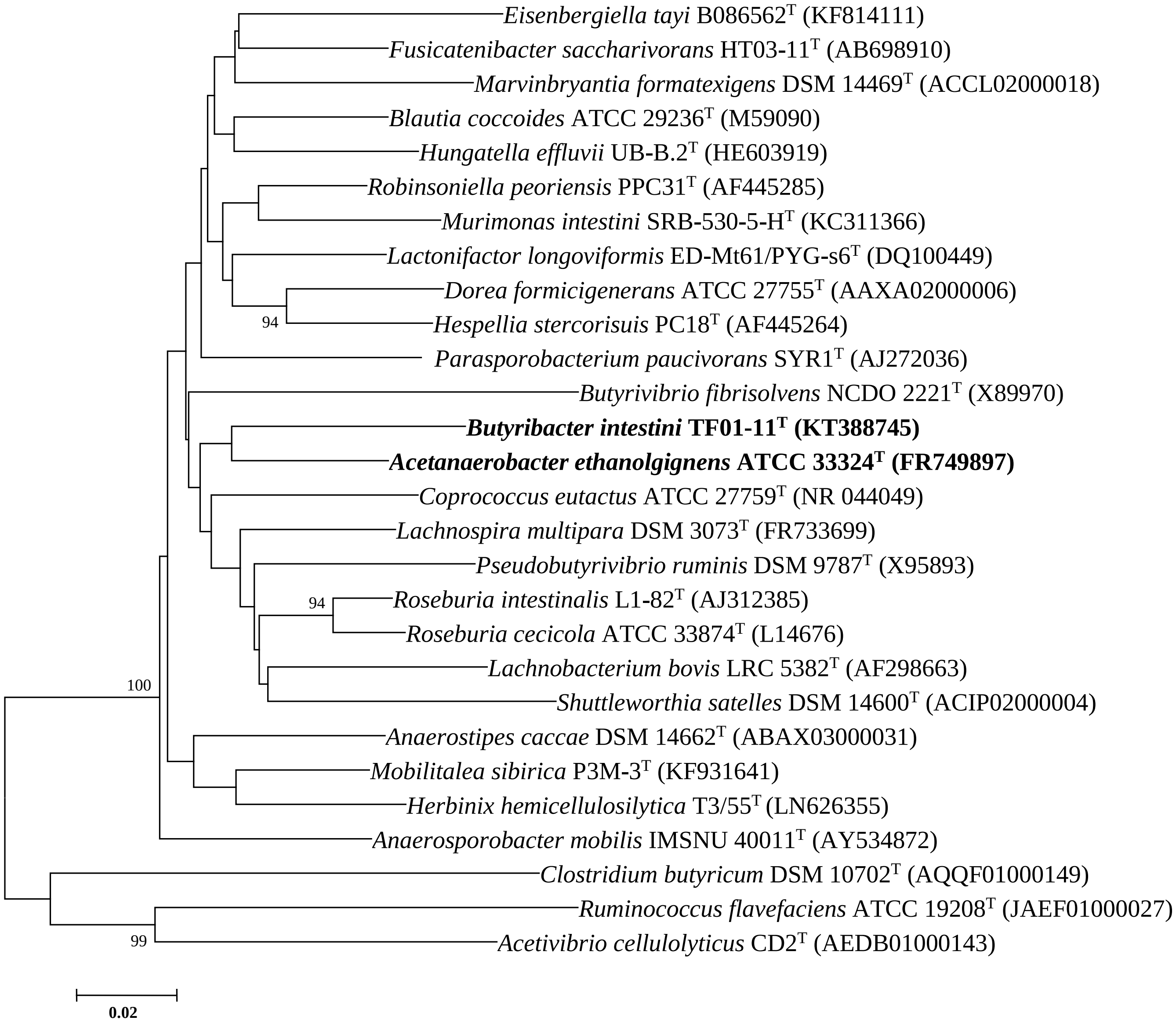
